# Putrescine acetyltransferase (PAT/SAT1) dependent GABA synthesis in astrocytes

**DOI:** 10.1101/2023.05.15.540086

**Authors:** Jiwoon Lim, Mridula Bhalla, Mingu Gordon Park, Wuhyun Koh, C. Justin Lee

## Abstract

GABA synthesis in astrocytes mediates tonic inhibition to regulate patho-physiological processes in various brain regions. Monoamine oxidase B (MAO-B) has been known to be the most important metabolic enzyme for synthesizing GABA from the putrescine degradation pathway. MAO-B converts N^1^-acetylputrescine to N^1^-acetyl-γ-aminobutyraldehyde and hydrogen peroxide (H_2_O_2_). Putrescine acetyltransferase (PAT), also known as spermidine and spermine N^1^-acetyltransferase 1 (SAT1), has been thought to be a feasible candidate enzyme for converting putrescine to N^1^-acetylputrescine. However, it has not been rigorously investigated or determined whether PAT/SAT1 contributes to GABA synthesis in astrocytes. To investigate the contribution of PAT/SAT1 to GABA synthesis in astrocytes, we conducted sniffer patch and whole-cell patch experiments with gene silencing of PAT/SAT1 by *Sat1* shRNA expression. Our results showed that the gene silencing of PAT/SAT1 significantly decreased the MAO-B-dependent GABA synthesis, which was induced by putrescine incubation, leading to decreased Ca^2+^-dependent release of GABA *in vitro*. Additionally, we found that, from the brain slice *ex vivo*, putrescine incubation induces tonic GABA inhibition in dentate gyrus granule cells, which can be inhibited by MAO-B inhibitor, selegiline. Consistent with our *in vitro* results, astrocytic gene silencing of PAT/SAT1 significantly reduced putrescine incubation-induced tonic GABA current, possibly by converting putrescine to N^1^-acetylputrescine, a substrate of MAO-B. Our findings emphasize a crucial role of PAT/SAT1 in MAO-B-dependent GABA synthesis in astrocytes.

## Introduction

γ-aminobutyric acid (GABA) is a crucial amino acid and one of the primary neurotransmitters in the central nervous system (CNS). Dysregulated GABA signaling has been linked to a range of brain diseases, such as depression, schizophrenia, and seizures (Kalueff and Nutt, 2007; Marques et al., 2021; Treiman, 2001). Tonic inhibition is one of the ways in which GABAA receptor-mediated inhibition occurs (Farrant and Nusser, 2005). In tonic inhibition, which is different from phasic inhibition, ambient GABA persistently activates extrasynaptic GABA A receptors, inducing GABA-mediated tonic conductance. Tonic activation of GABA A receptors regulates neuronal excitability, maintaining the balance between excitation and inhibition and filtering out irrelevant information. We have previously reported that astrocyte release GABA through Bestrophin1 to mediate tonic inhibition in diverse brain regions, including the cerebellum, hippocampus, thalamus, and substantia nigra pars compacta (Heo et al., 2020; Kwak et al., 2020; Lee et al., 2010). Astrocytes actively participate in signal transmission through tonic inhibition, highlighting their crucial role in the regulation of brain function. Thus, identifying the enzymes for GABA synthesis in astrocytes is critical for understanding role of astrocytes in the brain function.

GABA synthesis in astrocytes is different from neuronal GABA synthesis. In the neuron, glutamate is the only source for GABA synthesis, and decarboxylation by glutamic acid decarboxylase (GAD) produces GABA. In astrocytes, putrescine is another source of GABA synthesis. We previously have shown that increased putrescine induces GABA synthesis in astrocytes. Increased putrescine levels in the hippocampal astrocytes have been observed in Alzheimer’s disease, and elevated putrescine levels increase GABA synthesis in astrocytes and tonic GABA current in the hippocampus (Ju et al., 2022). There are two ways of putrescine to GABA synthesis in astrocytes, diamine oxidase (DAO)-dependent and MAO-B-dependent pathways (Kwak et al., 2020). In the DAO-dependent GABA synthesis, putrescine is directly oxidized to γ-aminobutyraldehyde by DAO, and then γ-aminobutyraldehyde is converted to GABA by aldehyde dehydrogenase 1 family, member A1 (ALDH1A1). In contrast to the DAO-dependent pathway, there are four steps in the MAO-B-dependent pathway. N^1^-acetylputrescine converted from putrescine is oxidized by MAO-B and becomes N^1^-acetyl-γ-aminobutyraldehyde. N^1^-acetyl-γ-aminobutyraldehyde is dehydrogenated to N^1^-acetyl-GABA by ALDH1A1 and further deacetylated to GABA by sirtuin 2 (Bhalla et al., 2023). Every step except the first step has been defined in the astrocytic MAO-B-dependent GABA synthesis. Although PAT/SAT1 has been proposed as the enzyme for putrescine to N^1^-acetylputrescine conversion, PAT/SAT1’s role in GABA synthesis of astrocytes has not been rigorously investigated yet (Petroff, 2002; Yoon and Lee, 2014).

In this study, we hypothesized that PAT/SAT1, well-known as polyamine acetyltransferase, is the astrocytic enzyme involved in putrescine-to-GABA synthesis. To investigate the role of PAT/SAT1 in astrocytic GABA synthesis, we conducted immunocytochemistry to measure putrescine-induced GABA levels after silencing PAT/SAT1 expression in cultured hippocampal astrocytes. We also performed two-cell sniffer patch experiments to examine GABA synthesis and release from cultured astrocytes. Furthermore, we established an *ex vivo* model of putrescine incubation to induce tonic GABA current. Using this model, we measured tonic GABA currents induced by putrescine incubation and confirmed our findings in brain slices by silencing the astrocyte-specific gene for PAT/SAT1. Our results demonstrate that PAT/SAT1 is involved in the putrescine-to-GABA synthesis pathway in astrocytes.

## Materials and Methods

### Animals

All C57BL/6 mice were group-housed in a temperature- and humidity-controlled environment with a 12 h light/dark cycle and had free access to food and water. All animal care and handling were approved by the Institutional Animal Care and Use Committee of the Institute for Basic Science (IBS-2020-005; Daejeon, Korea). For the slice patch, 8- to 20-week-old male C57BL/6 mice were used.

## Methods

### Cell culture

Primary hippocampal astrocytes were prepared from 1-day postnatal C57BL/6 mice as previously described (Woo et al., 2012). The cerebral cortex was dissected free of adherent meninges, minced, and dissociated into a single-cell suspension by trituration. Dissociated cells were plated onto plates coated with 0.1 mg/ml poly-D-lysine (Sigma). Cells were grown in Dulbecco’s modified Eagle’s medium (DMEM, Corning) supplemented with 4.5 g/L glucose, L-glutamine, sodium pyruvate, 10 % heat-inactivated horse serum, 10 % heat-inactivated fetal bovine serum, and 1000 units/ml of penicillin–streptomycin. Cultures were maintained at 37 °C in a humidified atmosphere containing 5 % CO_2_. Three days later, cells were vigorously washed with repeated pipetting using medium and the media was replaced to remove debris and other floating cell types.

### Quantitative real-time RT-PCR

Quantitative real-time RT-PCR was carried out using SYBR Green PCR Master Mix as described previously (Kwak et al., 2020). Briefly, reactions were performed in duplicates in a total volume of 10 ml containing 10 pM primer, 4 ml cDNA, and 5 ml power SYBR Green PCR Master Mix (Applied Biosystems). The mRNA level of each gene was normalized to that of *Gapdh* mRNA. Fold-change was calculated using the 2ΔΔCT method. The following sequences of primers were used for real-time RT-PCR. *Gapdh* forward: 5’ - ACC CAG AAG ACT GTG GAT GG -3’; *Gapdh* reverse: 5’ -CAC ATT GGG GGT AGG AAC AC-3’; *Sat1* forward: 5’ - GAC CCC TGA AGG ACA TAG CA-3’; *Sat1* reverse: 5’ - CCG AAG CAC CTC TTC TTT TG-3’.

### Sniffer patch from primary cultured astrocytes

Three days before, *Sat1* shRNA was transfected into primary hippocampal astrocytes. On the day of sniffer patch, HEK 293T cells expressing GABAc sensor were dissociated, triturated, added onto the cover glass with cultured astrocytes, and then allowed to settle for at least 1 h before sniffer patching. After HEK cells settled, cultured astrocytes were incubated with 5 μM Fura-2AM (mixed with 1ml of external solution containing 5 ml of 20 % pluronic acid, Invitrogen) for 40 min and washed at room temperature and subsequently transferred to a microscope stage. External solution contained 150 mM NaCl, 10 mM HEPES, 3 mM KCl, 2 mM CaCl_2_, 2 mM MgCl_2_, 5.5 mM glucose, pH adjusted to pH 7.3 and osmolality to 325 mOsmolkg^−1^. Intensity images of 510-nm wavelength were taken at 340-nm and 380-nm excitation wavelengths using iXon EMCCD (DV887 DCS-BV, ANDOR technology). The two resulting images were used for ratio calculations in Axon Imaging Workbench version 11.3 (Axon Instruments). To induce Ca^2+^-dependent astrocyte gliotransmission by activating PAR1 and TRPA1 receptors, respectively, TFLLR was either locally puffed (Kwak et al., 2020) or poked with a glass pipette (Oh et al., 2019) as previously. GABAc-mediated currents were recorded from HEK 293T cells under voltage clamp (Vh = -60 mV) using Axopatch 200A amplifier (Axon Instruments), acquired with pClamp 11.3. Recording electrodes (4–7 MΩ) were filled with 110 mM Cs-gluconate, 30 mM CsCl, 0.5 mM CaCl_2_, 10 mM HEPES, 4 mM Mg-ATP, 0.3 mM Na_3_-GTP and 10 mM BAPTA (pH adjusted to 7.3 with CsOH and osmolality adjusted to 290–310 mOsmkg^−1^ with sucrose). For simultaneous recordings, Imaging Workbench was synchronized with pClamp 11.3. To normalize differences in GABAc receptor-expression on the HEK 293T cells, 100 μM of GABA in the bath was applied to maximally activate the GABA_C_ receptors after current recording. Normalization was then accomplished by dividing the current induced by GABA released from astrocytes by the current induced by bath application of GABA.

### Immunocytochemistry of cultured hippocampal astrocytes

Cultured primary astrocytes were fixed in 4 % PFA in PBS for 15 mins and incubated for 1.5 h in a blocking solution (0.3 % Triton-X, 4 % normal serum in 0.1 M PBS) and then immunostained with a mixture of primary antibodies [Guineapig anti-GABA; AB175 (1:200) and chicken anti-GFAP; AB5541 (1:500)] in a blocking solution at 4 °C on a shaker overnight. After washing in PBS 3 times, samples were incubated with corresponding fluorescent secondary antibodies [Donkey anti-guineapig alexa 488; 706-475-148 (1:200) and donkey anti-chicken alexa 405; 703-475-155 (1:200)] for 2 h and then washed with PBS 3 times. Finally, samples were mounted with fluorescent mounting medium (Dako) and dried. A series of fluorescent images were obtained with a Zeiss LSM900 confocal microscope and processed for further analysis using ImageJ program. Any alterations in brightness or contrast were equally applied to the entire image set. Specificity of primary antibody and immunoreaction was confirmed by omitting primary antibodies or changing fluorescent probes of the secondary antibodies.

### Virus injection

Mice were anesthetized with vaporized isoflurane and placed into stereotaxic frames (Kopf). The scalp was incised, and a hole was drilled into the skull above the dentate gyrus (anterior/posterior, −1.5mm; medial/ lateral, −1.2 or +1.2mm from bregma, dorsal/ventral, −1.8mm from the brain surface). The virus was loaded into a glass needle and injected bilaterally into the dentate gyrus at a rate of 0.2 μl min^−1^ for 5 min (1 μl per each site) using a syringe pump (KD Scientific). Virus was generated from Institute for Basic science virus facility (IBS virus facility). AAV-GFAP-mCh, Lenti-pSico-Scrambled shRNA-GFP, and Lenti-pSico-PAT/SAT1 shRNA-GFP viruses were used in each experiment. Mice were used for patch-clamp 3 weeks after the virus injection.

### Preparation of brain slices

Mice were anesthetized with isoflurane and decapitated to remove the brain. The brains were sectioned in ice-cold slicing solution (234 mM sucrose, 2.5 mM KCl, 104 mM MgSO_4_, 1.25 mM NaH_2_PO_4_, 24 mM NaHCO_3_, 0.5 mM CaCl_2_-2H_2_O and 11 mM glucose). Horizontal slices (300 μm thick) were prepared with a vibrating-knife microtome Linear Slicer Pro7 (D.S.K, Japan). For stabilization, slices were incubated in room temperature for at least 1 h in a solution containing 124 mM NaCl, 3 mM KCl, 6.5 mM MgSO_4_, 1.25 mM NaH_2_PO_4_, 26 mM NaHCO_3_, 1 mM CaCl_2_-2H_2_O and 10 mM glucose, and simultaneously equilibrated with 95 % O_2_/5 % CO_2_ at 25°C. In some experiments, slices were incubated with blockers during stabilization for at least 2 h and slices were incubated with putrescine and selegiline one hour before recording.

### Tonic GABA recording

Prepared slices were transferred to a recording chamber that was continuously perfused with ACSF solution (flow rate, 2 ml min^− 1^). The slice chamber was mounted on the stage of an upright Olympus microscope and viewed with a 63x water immersion objective (0.90 numerical aperture) with infrared differential interference contrast optics. Cellular morphology was visualized by a charge-coupled device camera and Imaging Workbench software (INDEC BioSystems). Whole-cell recordings were made from granule cell somata located in the DG. The holding potential was - 70 mV. Pipettes (resistance 6-8 MΩ) were filled with internal solution [135 mM CsCl, 4 mM NaCl, 0.5 mM CaCl_2_, 10 mM HEPES, 5 mM EGTA, 2 mM Mg–ATP, 0.5 mM Na_2_–GTP, and 10 mM QX-314, pH adjusted to 7.2 with CsOH (osmolarity, 278 to 285 mOsmolkg^−1^)]. Baseline current was stabilized with d-AP5 (50 μM) and 6-cyano-7-nitroquinoxaline-2,3-dione (20 μM) before measuring tonic current. Electrical signals were digitized and sampled at 50 ms intervals with Digidata 1440 A and a MultiClamp 700B amplifier (Molecular Devices) using pCLAMP10.2 software. Data were filtered at 2 kHz. The amplitude of tonic GABA currents was measured by the baseline shift after bicuculline (100 μM) administration using the Clampfit program. Tonic current was measured from the baseline to bicuculline-treated current. Frequency and amplitude of spontaneous inhibitory postsynaptic currents before bicuculline administration were detected and measured by MiniAnalysis (Synaptosoft).

## Results

### Gene silencing of PAT/SAT1 decreases GABA synthesis and release in hippocampal astrocytes *in vitro*

We hypothesized that PAT/SAT1 has a role as the first enzyme in MAO-B-dependent GABA synthesis in astrocytes (Fig. 1A). To investigate this, we expressed *Sat1* shRNA in cultured hippocampal astrocytes *in vitro* to first determine whether *Sat1* shRNA reduces *Sat1* mRNA expression by quantitative RT-PCR (Fig. 1B). Compared to the control with scrambled shRNA, *Sat1* shRNA reduced the expression of *Sat1* mRNA by 91.61 %, showing that the gene silencing was effective. Next, we investigated whether *Sat1* shRNA reduces GABA synthesis and the amount of GABA in hippocampal astrocytes. Hippocampal astrocytes are known to have low amounts of basal putrescine and GABA under physiological conditions (Jo et al., 2014). It is noteworthy that PAT/SAT1 is involved in putrescine generation from spermidine and spermine degradation. To exclude the possibility that gene silencing of PAT/SAT1 decrease putrescine itself and examine the role of PAT/SAT1 in putrescine to GABA synthesis, hippocampal astrocytes were incubated with exogenous 180 μM putrescine. First, intracellular GABA contents in astrocytes were tested by immunocytochemistry. In the scrambled condition, putrescine incubation increased GABA intensity in astrocytes compared to the vehicle control (Fig. 1D, E). However, in the *Sat1* shRNA condition, the increase in GABA intensity by putrescine incubation was significantly reduced compared to the scrambled control (Fig. 1D, E). Interestingly, the increase in GABA with putrescine was accompanied by an increase in GFAP intensity (Fig. 1F), supporting the idea that H_2_O_2_, an additional product of MAO-B-dependent GABA synthesis, may induce astrocytic reactivity (Chun et al., 2020; Chun et al., 2022).

**Figure 1.**
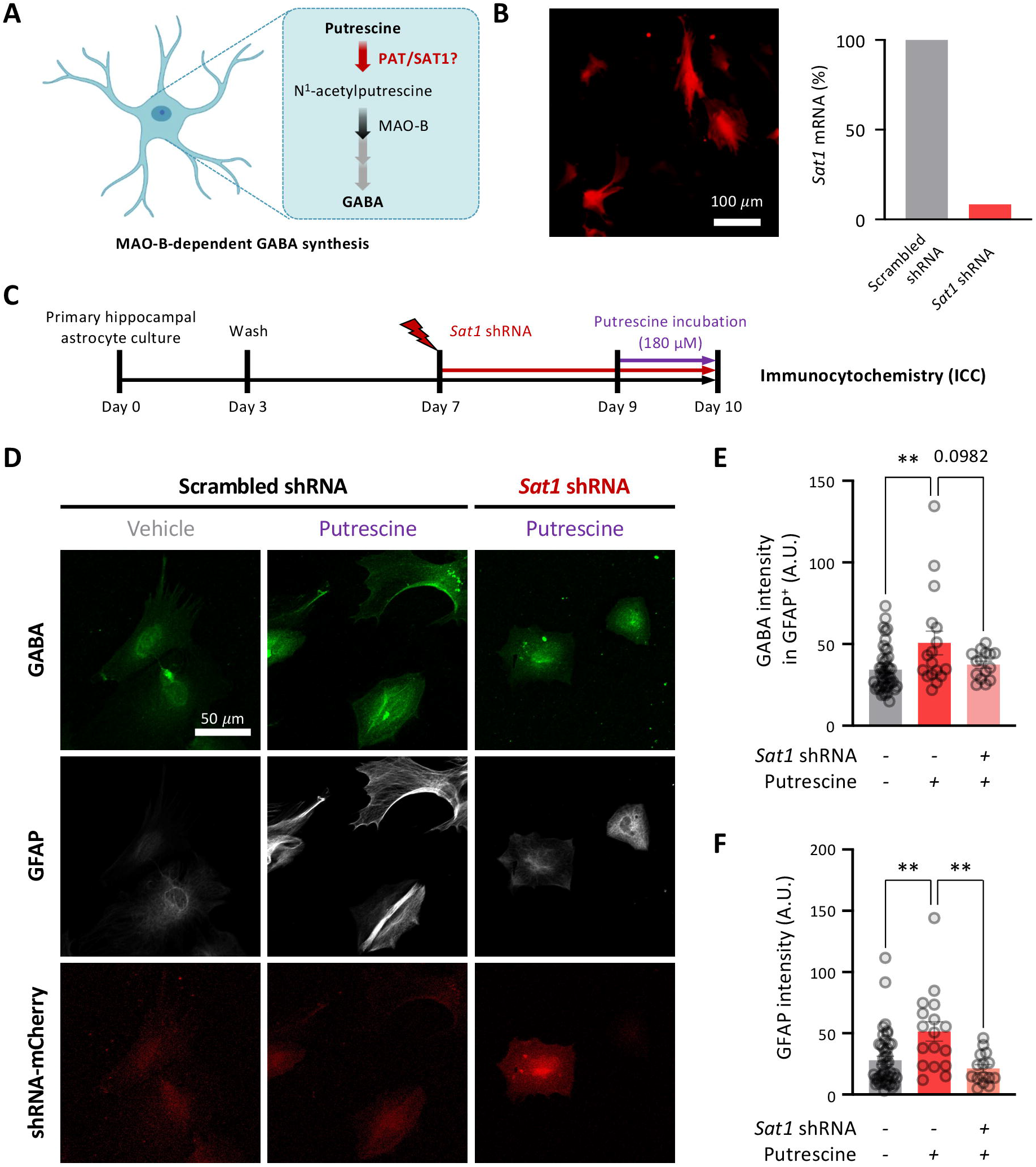
Putrescine acetyltransferase (PAT/SAT1) is a key enzyme in MAO-B-dependent GABA synthesis in hippocampal astrocytes A. Schematic diagram for MAO-B-dependent GABA synthesis in astrocytes. B. Representative image of *Sat1* shRNA transfected hippocampal astrocytes and *Sat1* mRNA level in shRNA transfected hippocampal astrocytes. C. Experimental timeline of immunocytochemistry. D. lmmunostaining of GABA and GFAP in shRNA transfected hippocampal astrocytes. E. GABA intensity in GFAP positive area. F. GFAP intensity. Data represent mean± SEM ** p<0.01 (One-way ANOVA).

We next attempted to determine if reduced GABA synthesis through gene silencing of PAT/SAT1 would also lead to reduced GABA release from hippocampal astrocytes (Fig. 2A). To measure GABA released by solitary hippocampal astrocytes, we performed the sniffer-patch technique as previously described (Jo *et al*., 2014; Kwak *et al*., 2020; Lee and Yoon, 2014) utilizing HEK293T cell that expresses GABAc receptors (referred to as a sensor cell). After seeding the hippocampal astrocyte with the sensor cell, we measured the amount of GABA released by the astrocyte by measuring the whole-cell patch current from the sensor cell (Fig. 2B, E). Given that cytosolic Ca^2+^ is a critical signal for GABA release in astrocytes, and that astrocytic Ca^2+^ can be elevated through either Ca^2+^ release from endoplasmic reticulum (ER) or Ca^2+^ influx from extracellular space, we investigated astrocytic GABA release with two methods to increase astrocytic Ca^2+^. First, to increase astrocytic Ca^2+^ through Ca^2+^ release from ER, TFLLR, a PAR1 agonist, was locally applied to hippocampal astrocyte (Fig. 2B-D). TFLLR application increased astrocytic Ca^2+^ in both scrambled and *Sat1* shRNA conditions, but GABA current from sensor cells was significantly reduced in *Sat1* shRNA (Fig. 2D), indicating that gene silencing of PAT/SAT1 decreased ER Ca^2+^-dependent GABA release from solitary astrocytes. Next, to increase astrocytic Ca^2+^ through Ca^2+^ influx from extracellular space, mechanical stimulation was delivered with microelectrode poking (Fig. 2E-G), activating astrocytic mechanosensitive channels (e.g. TRPA1) (Oh *et al*., 2019). Similar to TFLLR, poking stimulation elicited Ca^2+^ responses in both conditions, but GABA current from sensor cells was significantly reduced in *Sat1* shRNA (Fig. 2G). Taken together, these results indicate that PAT/SAT1 is necessary for MAO-B-dependent GABA synthesis and subsequent Ca^2+^-dependent GABA release in hippocampal astrocytes *in vitro*.

**Figure 2.**
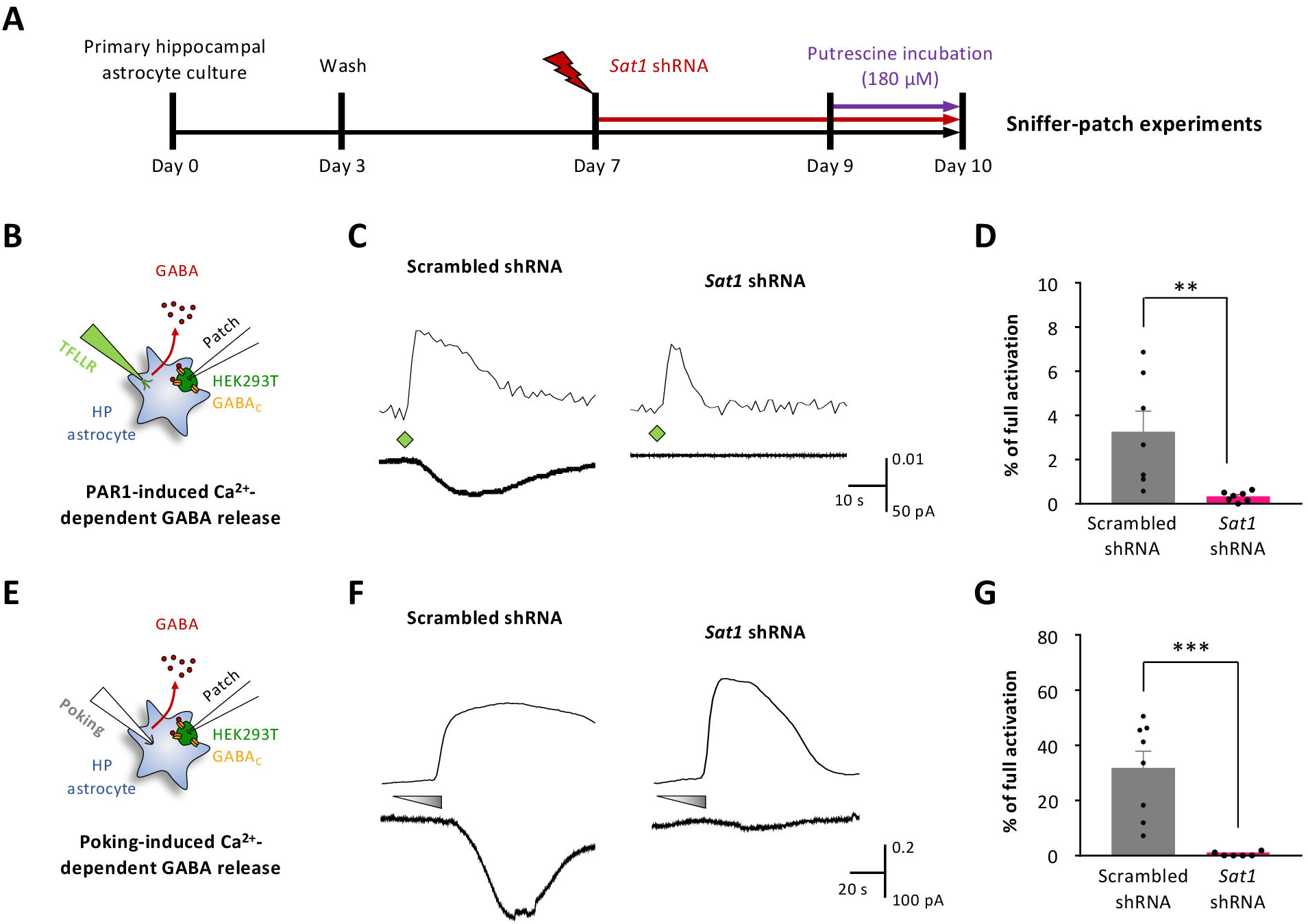
Knockdown of PAT/SAT1 decreases Ca^2+^-dependent GABA release from astrocyte *in vitro.* A. Experimental timeline for sniffer-patch. B. Experimental diagram for sniffer-patch of GABA release by PAR1 agonist. C. Representative traces of sensor current induced by GABA from primary cultured hippocampal astrocytes. D. Bar graph of sensor current. E. Experimental diagram for sniffer-patch of GABA release by mechanostimulation. F. Representative traces of sensor current induced by GABA from primary cultured hippocampal astrocytes. G. Bar graph of sensor current. Data represent mean ± SEM ** p<0.01, *** p<0.001 (Student’s t test).

### Astrocytic PAT/SAT1 is necessary for tonic GABA inhibition *ex vivo*

We showed that putrescine incubation induces GABA synthesis in the primary hippocampal astrocyte culture and GABA synthesis in astrocytes is PAT/SAT1-dependent in the culture system. However, we have not tested whether astrocytic PAT/SAT1 is involved in tonic GABA current in the hippocampus. In the physiological condition, tonic GABA current is not big enough to investigate the necessity of PAT/SAT1 in GABA synthesis. In addition, putrescine level would be reduced under physiological level by gene silencing of PAT/SAT1 which generate putrescine through spermidine and spermine degradation. So, we incubated brain slices with putrescine and conducted a whole-cell patch clamp to test whether putrescine incubation increases tonic GABA current in the hippocampus. To test whether putrescine incubation increases tonic GABA current and whether increased tonic GABA inhibition is dependent on the enzyme MAO-B, we conducted whole-cell patch clamp recordings of tonic GABA inhibition in granule cells of the dentate gyrus. We used the MAO-B inhibitor, selegiline, to ensure the specificity of the observed effects. To minimize variation among slices due to incubation time, we pre-incubated each slice with putrescine and selegiline separately for one hour at room temperature before recording (Fig. 3A). Our results showed that putrescine incubation significantly increased tonic GABA current, an effect that was blocked by selegiline (Fig. 3B, C). We did not observe a significant difference in the frequency of spontaneous inhibitory postsynaptic currents (sIPSCs), but unexpectedly the amplitude of sIPSCs was increased by putrescine incubation. Nevertheless, these findings indicate that putrescine increased tonic GABA inhibition through MAO-B-dependent GABA synthesis.

**Figure 3.**
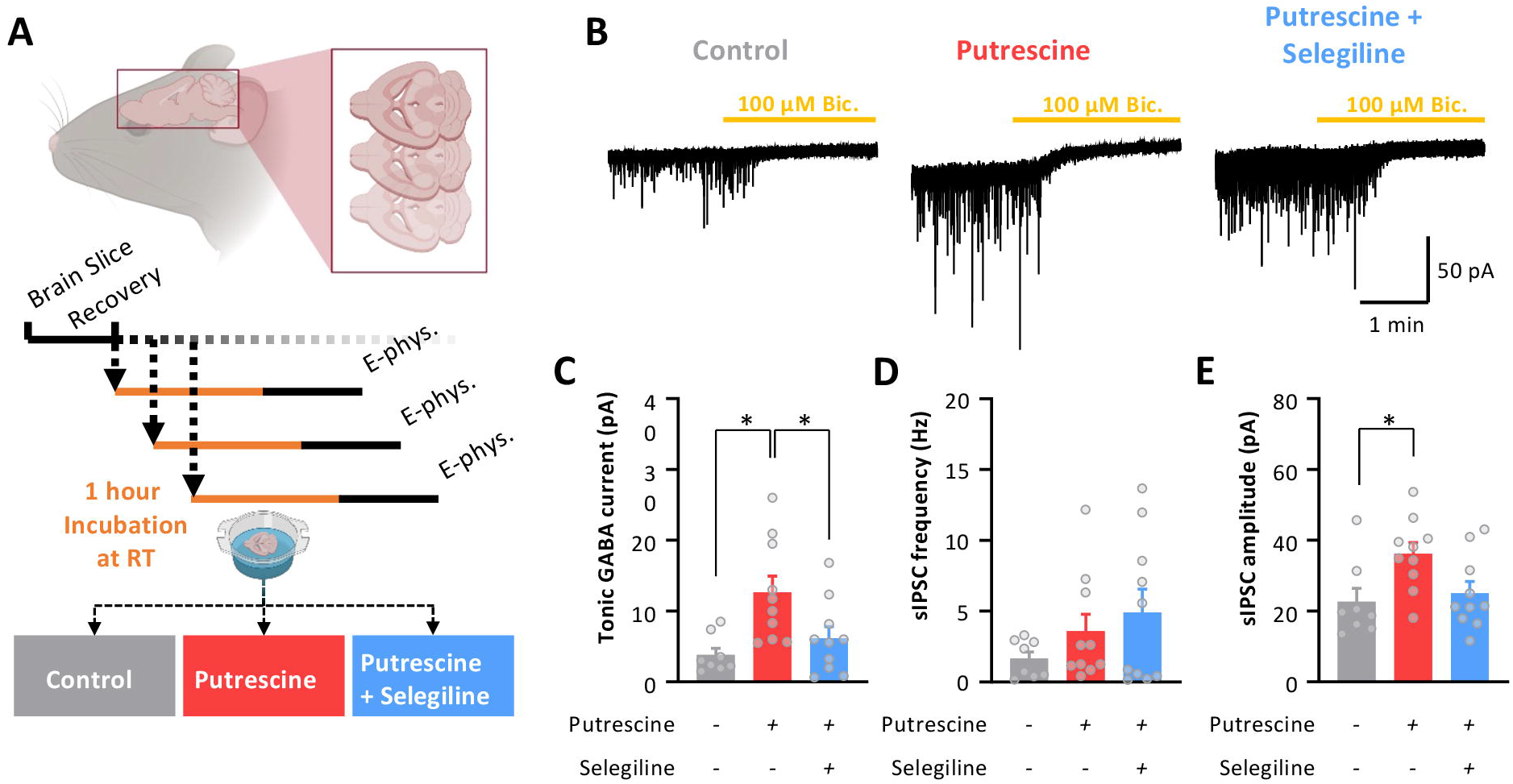
Increased tonic GABA inhibition by putrescine incubation A. Experimental timeline for tonic GABA recording B. Representative traces of GABAA receptor-mediated currents recorded from granule cells in the hippocampal slices incubated with putrescine or putrescine + selegilline. C. Bar graph of tonic GABA current amplitude. D. Bar graph of slPSC frequency before bicuculline treatment. E. Bar graph of slPSC amplitude before bicuculline treatment. Data represent mean± SEM * p<0.05 (One-way ANOVA).

To investigate whether astrocytic PAT/SAT1 is essential for tonic GABA inhibition induced by putrescine incubation, mice were injected with lentivirus carrying pSico-PAT shRNA-GFP which is Cre-dependent and AAV-GFAP-Cre-mCherry virus into the DG (Fig. 4A). More than 3 weeks after virus injection, we performed whole-cell patch clamp recordings in granule cells of DG (Fig. 4B). We observed that astrocyte-specific PAT/SAT1 gene silencing significantly reduced tonic GABA increased by putrescine incubation (Fig. 4C, D). Therefore, we concluded that astrocytic PAT/SAT1 is necessary for the tonic GABA release induced by putrescine incubation.

**Figure 4.**
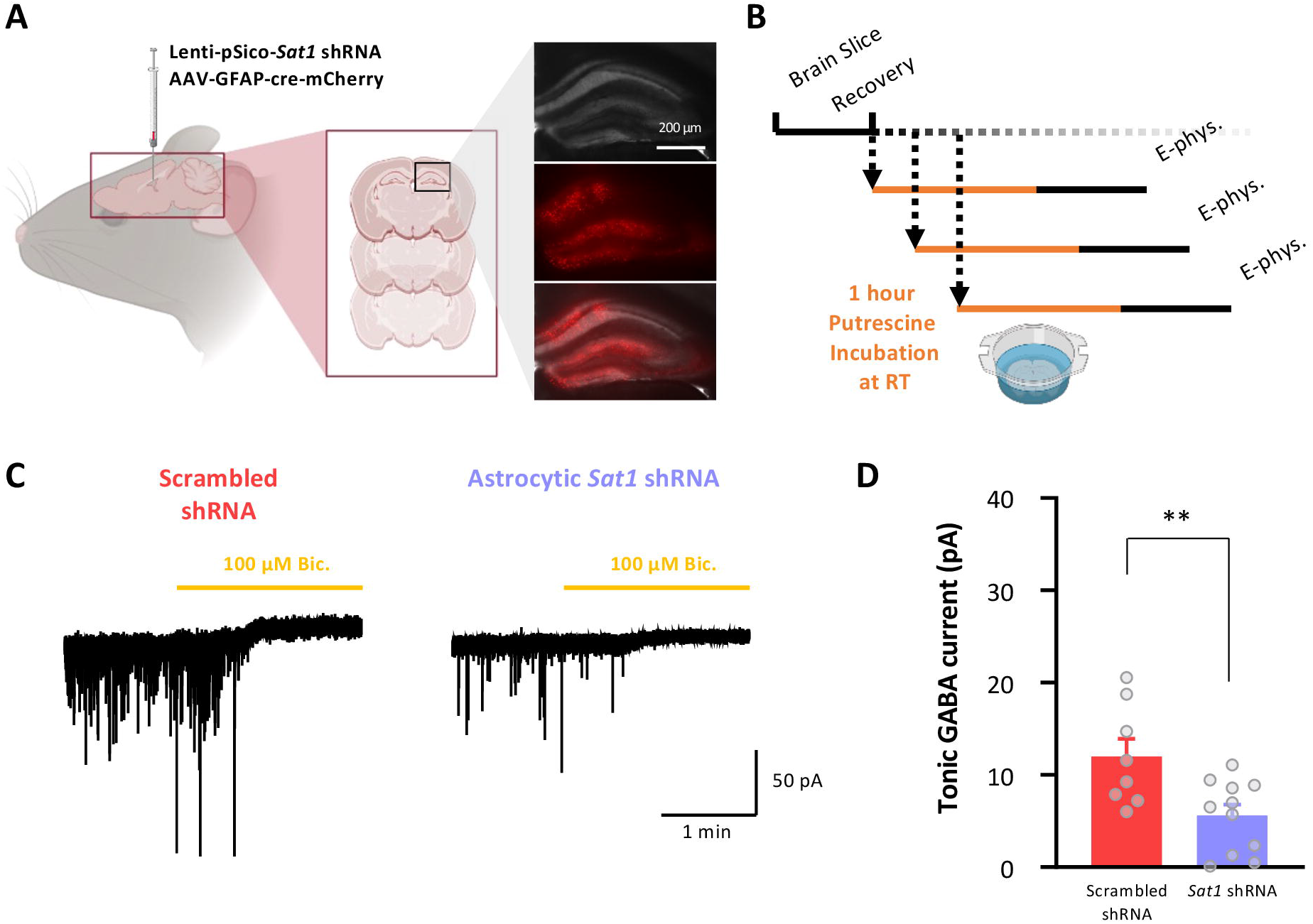
Knockdown of PAT/SAT1 decreases GABA release from astrocyte *ex vivo.* A. Experimental diagram and image of virus injection for astrocytic *Sat1* gene silencing. B. Experimental timeline for tonic GABA recording. C. Representative traces of GABAA receptor-mediated currents recorded from granule cells in the hippocampal slices incubated with putrescine. D. Bar graph of tonic GABA current amplitude. Data represent mean ± SEM ** p<0.01 (Student’s t test).

## Discussion

In this study, we have identified the enzyme responsible for the first step in astrocytic putrescine to GABA synthesis. We observed that putrescine-induced GABA synthesis and release is eliminated by gene silencing of PAT/SAT1 in primary-cultured mouse hippocampal astrocytes, as demonstrated by immunostaining (Fig. 1) and sniffer cell patch (Fig. 2). Moreover, we developed a putrescine incubation model to induce MAO-B-dependent tonic GABA in brain slices, which is typically undetectable under physiological conditions (Fig. 3). Using this newly developed model for tonic GABA induction, we observed that increased tonic GABA is reduced by astrocyte-specific gene silencing of PAT/SAT1 (Fig. 4). These findings demonstrate that PAT/SAT1 is the necessary enzyme for GABA synthesis in astrocytes, and possibly catalyzing the first step of MAO-B dependent GABA synthesis from putrescine. Metabolic analysis should be conducted to confirm the PAT/SAT1 activity of the conversion from putrescine to N^1^-acetylputrescine.

PAT/SAT1 has been well known as an acetyltransferase for degradation of polyamine including spermidine and spermine. Polyamines modulate the structure or activity of negatively charged molecules such as nucleic acids and proteins, and polyamine levels are tightly regulated. Dysregulated polyamines have been used as markers for disease, and several studies have focused on polyamine metabolism in astrocytes. Previous research has reported increased polyamine catabolism in astrocytes in various brain diseases and has explained the symptoms of the disease by loss of polyamine function (Merali et al., 2014; Sonninen et al., 2020). However, our results suggest that increased polyamine catabolism results in increased astrocytic MAO-B-dependent GABA synthesis. Our previous reports have shown that astrocytic MAO-B activation causes diverse diseases such as Alzheimer’s disease, Parkinson’s disease, traumatic brain injury and rheumatoid arthritis and symptoms of these diseases are alleviated by pharmacological or genetical inhibition of MAO-B (Chun et al., 2020; Chun et al., 2022; Heo et al., 2020; Jo et al., 2014; Park et al., 2019; Won et al., 2022). Therefore, excessive polyamine catabolism of astrocytes in pathological conditions results in not just loss of polyamine, but also aberrant H_2_O_2_ generation, the product of MAO-B-dependent polyamine degradation. Here, we showed that MAO-B-dependent GABA synthesis and release are blocked by astrocyte specific gene silencing of PAT/SAT1. Whether PAT/SAT1 inhibition can alleviate the symptoms of those diseases should be investigated in the future.

The acetylation of putrescine or spermidine and spermine by PAT/SAT1 activity is followed by oxidation through MAO-B or acetylpolyamine oxidase (APAO). During the oxidation of acetylpolyamine, H_2_O_2_, one of the reactive oxygen species, is generated (Casero and Pegg, 2009). Our research has shown that astrocytic MAO-B excessively produces H_2_O_2_ in pathological conditions (Chun et al., 2020; Chun et al., 2022). This excess H_2_O_2_ induces hypertrophy of reactive astrocytes and scar formation near injury sites in traumatic brain injury. Furthermore, H_2_O_2_generated by reactive astrocytes causes pathological symptoms such as tauopathy, neuronal death, and cognitive decline in Alzheimer’s disease, which can be alleviated by MAO-B inhibitor or H_2_O_2_scavenger (Chun et al., 2020). Therefore, inhibition of PAT/SAT1 activity would alleviate H_2_O_2_-derived symptoms by blocking not only MAO-B-dependent but also APAO-dependent H_2_O_2_ generation (Fig. 5). It would be interesting to determine the exact contribution of MAO-B and APAO in H_2_O_2_ generation in future study.

**Figure 5.**
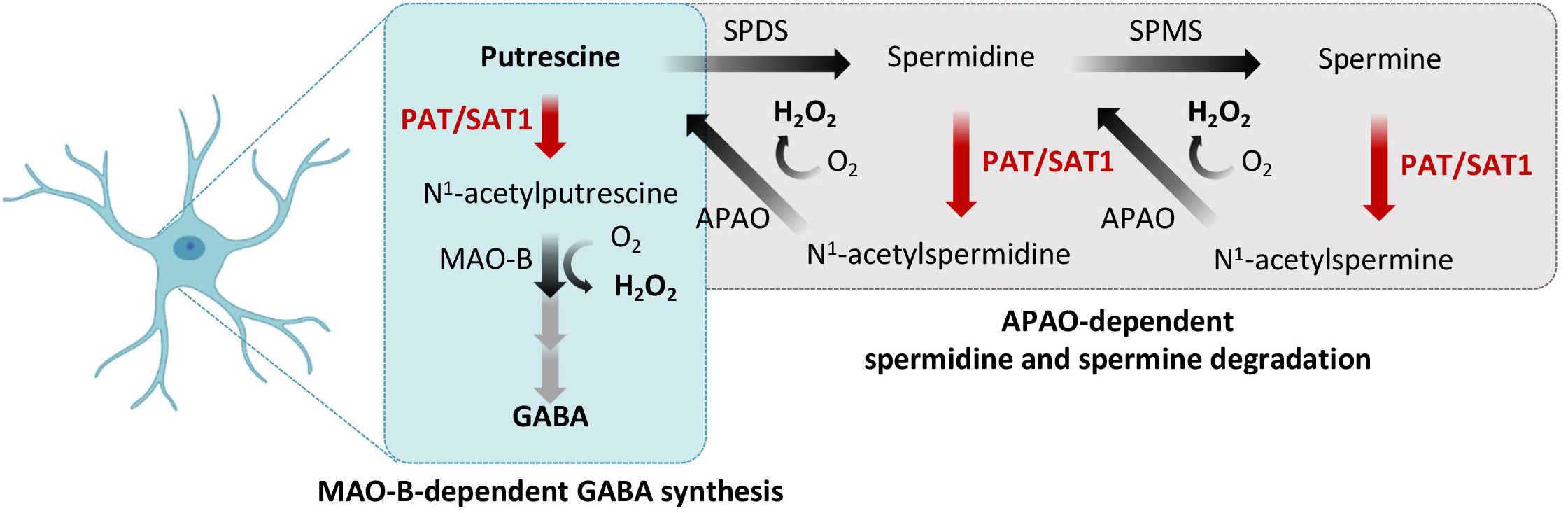
Schematic diagram of PAT/SAT1’s role in MAO-B-dependent GABA synthesis and APAO-dependent spermidine and spermine degradation PAT/SAT1: Putrescine acetyltransferase/Spermidine, spermine acetyltransferase, MAO-B: Monoamine oxidase B, GABA: γ-aminobutyric acid, SPDS: Spermidine synthase, SPMS: Spermine synthase, APAO: Acetylpolyamine oxidase

In addition to reducing harmful molecule, inhibiting PAT/SAT1 would increase spermidine and spermine, which have been shown to have neuroprotective roles in CNS (Frühauf et al., 2015; Ghosh et al., 2020; Liang et al., 2021; Schroeder et al., 2021; Xu et al., 2020; Yatin et al., 2001). There is a report that spermidine intake improves the cognitive function in the physiological conditions of drosophila, mice and humans (Schroeder *et al*., 2021). Spermidine and spermine have been shown to improve memory and delay aging in pathological conditions. There are several studies on the neuroprotective function of spermidine and spermine, including their ability to work as radical scavengers, maintain mitochondrial function and enhance autophagy (Frühauf *et al*., 2015; Ghosh *et al*., 2020; Liang *et al*., 2021; Xu *et al*., 2020; Yatin *et al*., 2001). Thus, PAT/SAT1 inhibition could induce accumulation of spermidine and spermine by reducing APAO-dependent degradation, which could ameliorate disease symptoms. Based on these reports, we propose that PAT/SAT1 is a potential therapeutic target for neurodegenerative diseases. Although there is a PAT/SAT1 inhibitor, diminazene aceturate, it is also known to activate angiotensin-converting enzyme 2 (ACE2), which is involved in the function of diverse organs such as the heart, kidney, and lung (Duan et al., 2020). Therefore, a PAT/SAT1-specific inhibitor should be developed to block H_2_O_2_ generation and enhance spermidine and spermine level without causing unintended side effects, enabling investigation of polyamine catabolism in the futures.

## Acknowledgments

This study was supported by the Institute for Basic Science (IBS), Center for Cognition and Sociality (IBS-R001-D2) to C.J.L., and Young Scientist Fellowship (IBS-R001-Y1) to W.K.

## References

Bhalla, M., Shin, J.I., Ju, Y.H., Park, Y.M., Yoo, S., Lee, H., and Lee, C.J.J. (2023). Molecular identification of ALDH1A1 and SIRT2 in the astrocytic putrescine-to-GABA metabolic pathway. bioRxiv, 2023.2001. 2011.523573.

Casero, Robert A., Jr, and Pegg, Anthony E. (2009). Polyamine catabolism and disease. Biochemical Journal 421, 323–338. 10.1042/bj20090598.

Chun, H., Im, H., Kang, Y.J., Kim, Y., Shin, J.H., Won, W., Lim, J., Ju, Y., Park, Y.M., and Kim, S. (2020). Severe reactive astrocytes precipitate pathological hallmarks of Alzheimer’s disease via H2O2– production. Nature neuroscience 23, 1555–1566.

Chun, H., Lim, J., Park, K.D., and Lee, C.J. (2022). Inhibition of monoamine oxidase B prevents reactive astrogliosis and scar formation in stab wound injury model. Glia 70, 354–367.

Duan, R., Xue, X., Zhang, Q.-Q., Wang, S.-Y., Gong, P.-Y., Yan, E., Jiang, T., and Zhang, Y.-D. (2020). ACE2 activator diminazene aceturate ameliorates Alzheimer’s disease-like neuropathology and rescues cognitive impairment in SAMP8 mice. Aging (Albany NY) 12, 14819.

Farrant, M., and Nusser, Z. (2005). Variations on an inhibitory theme: phasic and tonic activation of GABAA receptors. Nature Reviews Neuroscience 6, 215–229.

Frühauf, P.K.S., Porto Ineu, R., Tomazi, L., Duarte, T., Mello, C.F., and Rubin, M.A. (2015). Spermine reverses lipopolysaccharide-induced memory deficit in mice. Journal of neuroinflammation 12, 1–11.

Ghosh, I., Sankhe, R., Mudgal, J., Arora, D., and Nampoothiri, M. (2020). Spermidine, an autophagy inducer, as a therapeutic strategy in neurological disorders. Neuropeptides 83, 102083.

Heo, J.Y., Nam, M.H., Yoon, H.H., Kim, J., Hwang, Y.J., Won, W., Woo, D.H., Lee, J.A., Park, H.J., Jo, S., et al. (2020). Aberrant Tonic Inhibition of Dopaminergic Neuronal Activity Causes Motor Symptoms in Animal Models of Parkinson’s Disease. Curr Biol 30, 276–291 e279. 10.1016/j.cub.2019.11.079.

Jo, S., Yarishkin, O., Hwang, Y.J., Chun, Y.E., Park, M., Woo, D.H., Bae, J.Y., Kim, T., Lee, J., Chun, H., et al. (2014). GABA from reactive astrocytes impairs memory in mouse models of Alzheimer’s disease. Nat Med 20, 886–896. 10.1038/nm.3639.

Ju, Y.H., Bhalla, M., Hyeon, S.J., Oh, J.E., Yoo, S., Chae, U., Kwon, J., Koh, W., Lim, J., and Park, Y.M. (2022). Astrocytic urea cycle detoxifies Aβ-derived ammonia while impairing memory in Alzheimer’s disease. Cell Metabolism 34, 1104–1120. e1108.

Kalueff, A.V., and Nutt, D.J. (2007). Role of GABA in anxiety and depression. Depression and anxiety 24, 495–517.

Kwak, H., Koh, W., Kim, S., Song, K., Shin, J.-I., Lee, J.M., Lee, E.H., Bae, J.Y., Ha, G.E., and Oh, J.-E. (2020). Astrocytes control sensory acuity via tonic inhibition in the thalamus. Neuron 108, 691–706. e610.

Lee, C.J., and Yoon, B.-E. (2014). Protease-activated receptor 1-induced GABA release in cultured cortical astrocytes pretreated with GABA is mediated by the Bestrophin-1 channel. Animal Cells and Systems 18, 244–249.

Lee, S., Yoon, B.-E., Berglund, K., Oh, S.-J., Park, H., Shin, H.-S., Augustine, G.J., and Lee, C.J. (2010). Channel-mediated tonic GABA release from glia. Science 330, 790–796.

Liang, Y., Piao, C., Beuschel, C.B., Toppe, D., Kollipara, L., Bogdanow, B., Maglione, M., Lützkendorf, J., See, J.C.K., and Huang, S. (2021). eIF5A hypusination, boosted by dietary spermidine, protects from premature brain aging and mitochondrial dysfunction. Cell reports 35, 108941.

Marques, T.R., Ashok, A.H., Angelescu, I., Borgan, F., Myers, J., Lingford-Hughes, A., Nutt, D.J., Veronese, M., Turkheimer, F.E., and Howes, O.D. (2021). GABA-A receptor differences in schizophrenia: a positron emission tomography study using [11C]Ro154513. Molecular Psychiatry 26, 2616–2625. 10.1038/s41380-020-0711-y.

Merali, S., Barrero, C.A., Sacktor, N.C., Haughey, N.J., Datta, P.K., Langford, D., and Khalili, K. (2014). Polyamines: predictive biomarker for HIV-associated neurocognitive disorders. Journal of AIDS & clinical research 5, 1000312.

Oh, S.J., Lee, J.M., Kim, H.B., Lee, J., Han, S., Bae, J.Y., Hong, G.S., Koh, W., Kwon, J., Hwang, E.S., et al. (2019). Ultrasonic Neuromodulation via Astrocytic TRPA1. Curr Biol 29, 3386–3401 e3388. 10.1016/j.cub.2019.08.021.

Park, J.-H., Ju, Y.H., Choi, J.W., Song, H.J., Jang, B.K., Woo, J., Chun, H., Kim, H.J., Shin, S.J., and Yarishkin, O. (2019). Newly developed reversible MAO-B inhibitor circumvents the shortcomings of irreversible inhibitors in Alzheimer’s disease. Science Advances 5, eaav0316.

Petroff, O.A. (2002). Book review: GABA and glutamate in the human brain. The Neuroscientist 8, 562–573.

Schroeder, S., Hofer, S.J., Zimmermann, A., Pechlaner, R., Dammbrueck, C., Pendl, T., Marcello, G.M., Pogatschnigg, V., Bergmann, M., and Müller, M. (2021). Dietary spermidine improves cognitive function. Cell reports 35, 108985.

Sonninen, T.-M., Hämäläinen, R.H., Koskuvi, M., Oksanen, M., Shakirzyanova, A., Wojciechowski, S., Puttonen, K., Naumenko, N., Goldsteins, G., and Laham-Karam, N. (2020). Metabolic alterations in Parkinson’s disease astrocytes. Scientific reports 10, 14474.

Treiman, D.M. (2001). GABAergic mechanisms in epilepsy. Epilepsia 42, 8–12.

Won, W., Choi, H.-J., Yoo, J.-Y., Kim, D., Kim, T.Y., Ju, Y., Park, K.D., Lee, H., Jung, S.Y., and Lee, C.J. (2022). Inhibiting peripheral and central MAO-B ameliorates joint inflammation and cognitive impairment in rheumatoid arthritis. Experimental & Molecular Medicine 54, 1188–1200.

Woo, D.H., Han, K.-S., Shim, J.W., Yoon, B.-E., Kim, E., Bae, J.Y., Oh, S.-J., Hwang, E.M., Marmorstein, A.D., and Bae, Y.C. (2012). TREK-1 and Best1 channels mediate fast and slow glutamate release in astrocytes upon GPCR activation. Cell 151, 25–40.

Xu, T.-T., Li, H., Dai, Z., Lau, G.K., Li, B.-Y., Zhu, W.-L., Liu, X.-Q., Liu, H.-F., Cai, W.-W., and Huang, S.-Q. (2020). Spermidine and spermine delay brain aging by inducing autophagy in SAMP8 mice. Aging (Albany NY) 12, 6401.

Yatin, S.M., Yatin, M., Varadarajan, S., Ain, K.B., and Butterfield, D.A. (2001). Role of spermine in amyloid β-peptide-associated free radical-induced neurotoxicity. Journal of neuroscience research 63, 395–401.

Yoon, B.-E., and Lee, C.J. (2014). GABA as a rising gliotransmitter. Frontiers in neural circuits 8, 141.

